# Female genetic variation controlling timing of mating plug ejection in *Drosophila melanogaster*

**DOI:** 10.64898/2026.06.27.734984

**Authors:** Jolie A. Carlisle, Rachel M. J. Craig, Mikaela Matera-Vatnick, Bianca M. Villanueva, Adriana R. Andrus, Elissa J. Cosgrove, Dawn S. Chen, Andrew G. Clark, Mariana F. Wolfner

## Abstract

In multiply-mating species, male-female postcopulatory, prezygotic interactions can influence reproductive outcomes. In *Drosophila melanogaster*, females can bias sperm storage and usage and thereby influence paternity outcomes. One mechanism by which females may regulate paternity contributions from specific males is through modulation of mating plug ejection timing. The *D. melanogaster* mating plug is composed of seminal fluid proteins, and some female-derived proteins, that coagulate in the female reproductive tract during mating. The mating plug facilitates sperm storage; thus, timing of female mating plug ejection is associated with sperm storage and relative paternity contributions in cases of multiple mating. However, whether there is natural genetic variation among females that shapes mating plug ejection timing, and genes or phenomena that might mediate it are unknown. We examined mating plug ejection in females from 69 lines of the *Drosophila* Genetic Reference Panel and observed dramatic differences in median plug ejection timing ranging from less than 1 to over 6 hours. We used this variation to perform a genome-wide association study to identify gene candidates associated with this phenotype. Many gene candidates are expressed in the brain and/or function in neurodevelopment. The candidate pool was also enriched for genes expressed in the ovary and functioning in oogenesis, indicating a link between female reproductive physiology and mating plug ejection. Consistent with this interpretation, females without a germline delay mating plug ejection. Our results demonstrate that female mating plug ejection is a physiologically integrated reproductive trait with a genetic basis that can be shaped by selection.

**Article Summary:** The *D. melanogaster* mating plug is composed of seminal fluid proteins and some female-derived proteins that coagulate in the female reproductive tract during mating. The mating plug facilitates sperm storage; thus, timing of female mating plug ejection is associated with sperm storage and relative paternity contributions in cases of multiple mating. Using the DGRP, we observed heritable genetic variation in female timing of mating plug ejection and through a GWAS find associated gene candidates. Gene candidates are enriched for neurodevelopment function and oogenesis function. We experimentally validate the connection between female mating plug ejection and the ovary.

## Introduction

Across species, males and females show selectivity in mate choice^1–5^. Given the energetic costs of courtship, copulation, and reproduction, as well as constraints on gamete production, preferentially mating with higher-quality partners can enhance reproductive success^6,7^. However, mate choice does not stop at copulation. In multiply mating species, both males and females may influence reproductive outcomes through post-copulatory, prezygotic interactions^8–12^. In internally fertilizing species, males may modulate reproductive investments through differential allocations of the quality or quantity of sperm or ejaculate molecules^13–15^. Females have been shown to molecularly or behaviorally control fertilization success from matings^16–19^. For either sex, this modulation allows strategic allocation of reproductive resources (e.g., gametes and energy) toward preferred partners. However, the molecular mechanisms that mediate these post-copulatory processes, also known as cryptic choice, are poorly understood in either sex in any species.

In *Drosophila*, both males and females may strategically influence post-mating, pre-zygotic interactions around the ejaculate^1,19–22^. Males have been demonstrated to adjust ejaculate investment in response to female quality and environmental cues such as the presence of rival males^14,15,23,24^. Females can bias sperm usage toward preferred males and influence sperm competition outcomes^9,25,26^. One potential mechanism that both males and females may exert influence involves controlling the time of expulsion of the mating plug – which contains the transferred sperm. During copulation, *Drosophila* males transfer seminal fluid in addition to sperm to their mates^27^. A subset of the proteins in the seminal fluid (Sfps, seminal fluid proteins) along with some female proteins coagulate to form the mating plug in the female uterus^28–31^. Disruption of mating plug formation causes reduced sperm storage due to ejaculate loss^28,31,32^. Furthermore, timing of mating plug ejection can influence number of sperm stored. Therefore, by controlling plug ejection, multiply mating females may bias sperm storage and usage^28,29,33–35^. The *Drosophila* plug also acts as a temporary chemical block to remating by carrying male-derived anti-aphrodisiac pheromone (cVA) which makes females less attractive, therefore, early ejection can lead to remating earlier ^36^. Therefore, since mating plug ejection timing affects both sperm storage and remating, both sexes may benefit from influencing this process to exert control over it

Recent work has begun to elucidate mechanisms of female control of mating plug ejection timing. Females lacking a spermathecae (a secretory and sperm storing organ of the female reproductive tract) fail to eject the mating plug, suggesting spermathecae-derived female proteins assist in efficient plug expulsion^30^. Several studies have examined neuronal mechanisms underlying female control of mating plug ejection timing. In particular, disruption of the neuropeptide Dh44 and its receptor accelerates female ejection of the mating plug^37^. Other studies have examined how signaling between the male and female can affect female neuronal control of mating plug ejection timing. *Drosophila melanogaster* females eject their mating plug more rapidly when in the presence of another male or a mated female, due to female detection of male pheromones accelerating mating plug ejection through a subset of pC1 neurons^21,38^. These findings provide insight into ways in which females can exert cryptic choice through regulation of mating plug ejection timing; however, the natural variation that shapes the phenotype and the genetic architecture that underlies it is unknown. To address this gap, we used the *Drosophila* Genetic Reference Panel (DGRP) to quantify natural genetic variation in female mating plug ejection timing when mated to a standard male, and to identify genes associated with this variation^39^. This unbiased genome-wide approach enables the identification of candidate mechanisms underlying female control of mating plug ejection timing.

We have observed that there is much heritable genetic variation contributing to differences in female mating plug ejection timing among DGRP lines, and our GWAS identified high-confidence gene candidates associated with this phenotype. We observed that many of the genes associated with these SNPs are linked to developmental functions, many of which are not yet characterized. However, digging deeper on shared phenotypic effects of mutant alleles of these gene candidates revealed enrichment for neurodevelopment and oogenesis phenotypes. We hypothesized that there is a link between the ovary function or physiology to female mating plug ejection, which we validated by observing that females without a germline show slower ejection of the mating plug. Together, our results demonstrate that female mating plug ejection is a physiologically integrated reproductive trait with a genetic basis that can be shaped by selection.

## Methods

### Fly stocks and rearing

Fly stocks were raised on standard glucose-yeast-agar media in a 12-hour light/dark cycle at 25°C. Standard Canton-S wild-type fly stocks were obtained from the Wolfner lab stocks. DGRP stocks were obtained from the Clark lab stocks and are available at the Bloomington Drosophila Stock Center. DGRP lines used can be found in Supplementary Dataset 2. To investigate mating plug ejection timing in females lacking a germline, we used females that were daughters of *tudor*^1^ homozygous mothers which were generated from a *tud*^1^ *bw*^1^*, sp*^1^/*SM5* stock provided to us from the lab of Dr. Bob Boswell^40^.

### Measurement of female mating plug ejection time

Flies were grown on standard media for mating plug ejection assays; after 7 days, the parents were removed, and unmated males and females were collected after eclosion under CO_2_ anesthesia. These unmated males and females were aged 3-5 days for assays in single-sex vials containing standard media. For all experiments, Canton-S wildtype males were used. Pairwise matings were set up in individual numbered vials containing standard media. The pairs were observed, and the time of introduction, start and end of mating were recorded. After copulation, females were transferred by aspiration to an individually numbered observation chamber for observation, and males were discarded. These small, black, 3D-printed observation chambers, whose openings are covered with a glass coverslip^41^, allowed observation of individual females for presence of the mating plug, which is auto-fluorescent under blue light. Females in chambers were checked every 10 minutes for the presence of a plug using a NightSEA fluorescent light system (SFA-RB). Once the plug was no longer observed within the female, the time was noted as the mating plug ejection time. If the female did not eject the mating plug during the five-hour observation window, the observation was ended and the female was marked as not having ejected. Supplementary Figure 1 shows a schematic of the experimental setup.

### Genome Wide Association Study of female mating plug ejection timing

Females from sixty-nine randomly selected DGRP lines were crossed to standard Canton-S males and used to conduct the GWAS^39,42^. A single male genotype was chosen to control for male-associated variation in order to increase power to detect loci associated with female variation. Also, within-line matings could mask variation obscured by male coadaptation with the female that may mask female variation. While this design increases power for detecting genes associated with female variation, it would not distinguish between loci impacting female-intrinsic variation vs variation due to male-female interaction. Therefore, some reported variation may reflect differences in female response to the single male genotype we used. For each experimental day approximately 50 pairwise matings were set up, with ∼20-50 individuals assayed. Six lines were assayed twice to determine the consistency of phenotypic estimation (Pearson’s correlation of 98.6%) (Supplementary Figure 7).

To account for right-censoring, because not all females had ejected the plug within the observation period after transfer to the ejection chambers, we used survival analysis to estimate the median mating plug ejection timing for each DGRP line on each experimental day. Specifically, we used last observation time for individuals that did not eject and fit a parametric survival regression model (R function survival::survreg(), log-normal distribution) to the data for a given DGRP line experiment. We used the log-transformed predicted median value from the fitted survival model for each experiment (line-day) as the GWAS phenotype. To test for genome-wide associations, we modeled mating plug ejection as a function of genotype, time, inversions, and Wolbachia infection, while accounting for random variability due to relatedness, replicate effects, and researcher differences (R function lme4qtl::relmatLmer()), using the formula: MPE ∼ gt + time + In.2L.t + In.2R.NS + In.3R.K + In.3R.Mo + Wolbachia + (1 | line) + (1 | line.ID) + (1 | researcher)) (Supplementary Dataset 2) (acronyms: mating plug ejection (MPE), genotype (gt), inversion (In)). Genotype, inversion status, Wolbachia infection status, were all obtained from associated data of the DGRP2 release and were downloaded from the DGRP2 website (http://dgrp2.gnets.ncsu.edu/)^42^. We included the count of observations for each experiment as the weights argument and used the genetic relatedness matrix (GRM) as the covariance structure for “line.ID” in the relmat argument (with “line” reflecting repeated measures). This model was compared to the same model without genotype by using a likelihood ratio test (LRT), and SNPs were ranked by LRT *P*-value. We used the DGRP2 file “dgrp.fb557.annot.txt” (from dgrp2.gnets.ncsu.edu/) for annotation of SNPs to genes.

To generate hypotheses from the results of the GWAS, we used Gene Ontology^43^, network STRING^44^ analysis, FlyAtlas2^45^ tissue-expression data, and Flybase^46^ annotations to look for trends in our gene candidate list. For more details on our approach, please see Supplementary Methods 1.

### Mating plug ejection by females without a germline

Flies lacking germline cells were generated by crossing unmated *tud*^1^ *bw*^1^ *sp*^1^ females to CS males. Genetically matched control flies with a germline were generated by crossing unmated *tud*^1^ *bw*^1^ *sp*^1^*/SM5* females to CS males and taking Cy+ progeny. Absence or presence of a germline was confirmed of experimental and control females by dissection to examine ovaries and ability to lay eggs. Pairwise matings of CS males mated to control or experimental females were assayed for mating plug ejection timing. Three independent biological replicates were performed. Mating plug ejection times for experimental and control females mated to Canton-S males were compared using a two-sided Welch’s *t*-test. To evaluate the overall effects across replicates, data were analyzed using a linear mixed-effects model (Value ∼ Treatment + (1|Replicate)), with replicate included as a random effect to account for baseline differences between replicates. Mixed-effects models were fitted using the lme4 R package with associated *P*-values calculated using the lmerTest package^47,48^.

Not all female flies ejected the mating plug within the five-hour observation window. We evaluated whether there was a significant difference in proportion of females ejecting the mating plug in the control vs. experimental group using a Fisher’s exact test with a significance threshold of *P* < 0.05. Odds ratios from the three biological replicates were combined using REML implemented in the metafor package in R and evaluated for heterogeneity^49^.

## Results

### Variation of female mating plug ejection timing in the DGRP

We investigated the genetic basis of female variation in mating plug ejection timing using over 3000 matings spread over 69 lines of the DGRP (Supplementary Dataset 1). We used the same male genotype across all experiments, in order to specifically investigate genetic variation contributed by the female. However, since only one male genotype was used, observed variation that was female intrinsic or based on male-female interaction were indistinguishable. Abundant variation was observed between DGRP lines for estimated timing of mating plug ejection (Figure 1, Supplementary Dataset 1 & 2). Median female mating plug ejection time varies among lines from under one hour to over 6 hours. The average of the estimated median mating plug ejection times for each line (and line replicates) mating plug was 2.81 hours ± 1.5 hours. Broad-sense heritability was estimated as the variance among lines in median ejection time from the survival-analysis (1.671) divided by the variance among all flies tested based on a survival model without line information (4.959), yielding 33.7%, indicating a moderate genetic contribution to the variability observed among lines. We investigated mating plug ejection timing on two separate days for 6 lines to estimate repeatability of the measurements (Supplementary Figure 7). Spearman’s rank correlation of line medians between the two measurement days was perfect (rho=1, p=.0028), indicating consistent ordering of genetic backgrounds. A strong Pearson correlation was observed between replicates (*r* = 0.986*, t* = 11.741, df = 4, *P*-value = 0.0003) indicating concordance in genotype-specific effect sizes across replicates.

**Figure 1.**
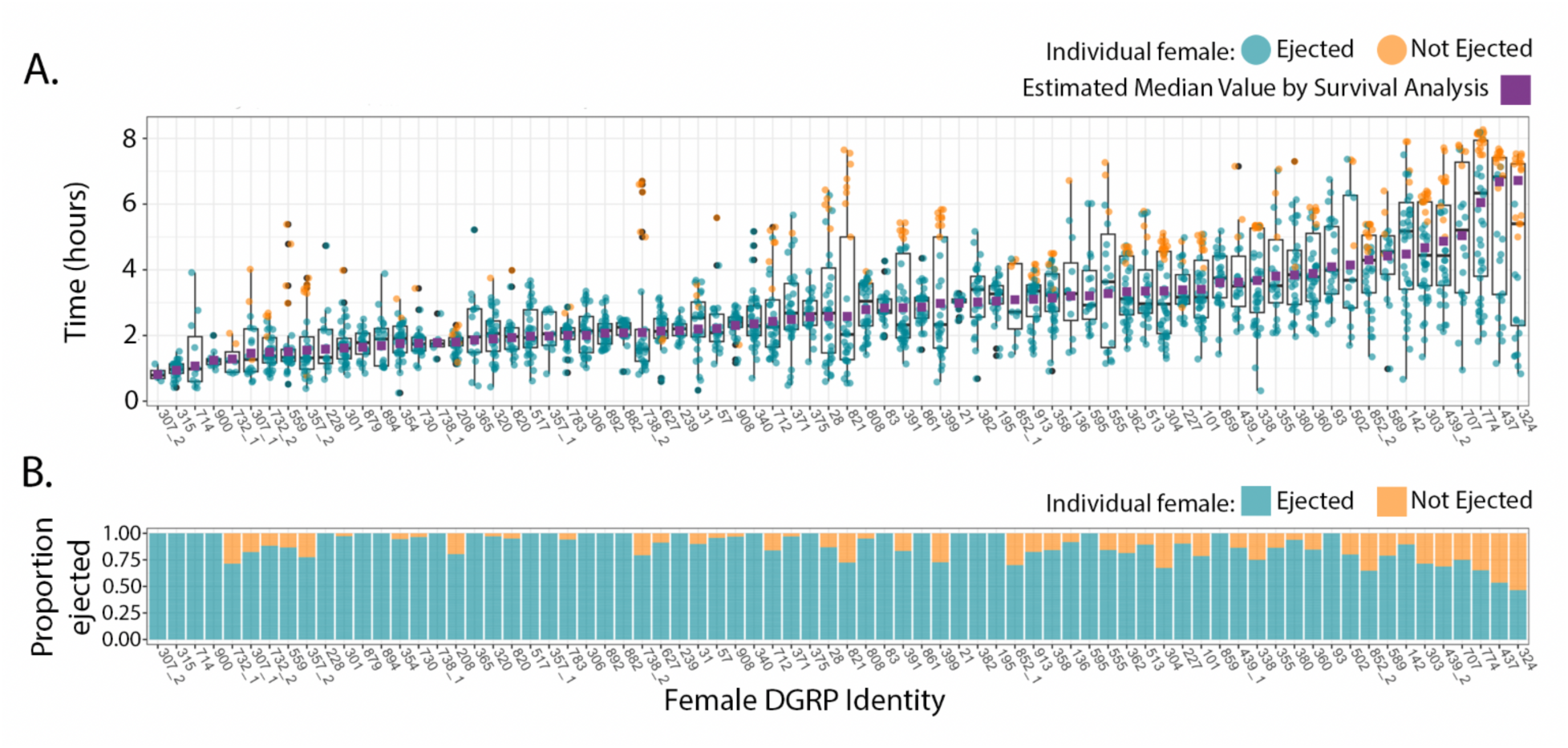
Variation in female mating plug ejection timing in 69 lines of the DGRP. A) Variation in mating plug ejection time across lines. Each point indicates the time from the end of mating to ejection (aquamarine) or the time from the end of mating until the last observation (orange) for flies that did not eject during the duration of the experiment. For the genome-wide association study, median mating plug ejection time estimated by survival analysis for each DGRP line (including replicates) was used and is indicated by a purple square. B) Proportion of mated females that ejected the mating plug within five hours after mating for each line and line replicate. Aquamarine indicates having ejected the mating plug during the observation window, and orange indicates not having ejected. Note that the DGRP lines are in the same order along the *x*-axis in both panels. Raw data available in Supplementary Dataset 1; Estimated median values are available in Supplementary Dataset 2.

### Significant SNPs associated with female mating plug ejection timing

To develop an understanding of the genetic basis of the extensive population-level variation observed for female mating plug ejection timing, we performed a genome-wide association study (Figure 2, Supplementary Dataset 4). The use of a single male genotype in data collection increased the power to detect SNPs based on female variation. For each line, we used the median mating plug ejection time estimated from survival analysis as the phenotype (two independent mean values were used for lines measured on two days) (Supplementary Dataset 2). We used a *P*-value cut-off of 1 x 10^-5^ yielding 43 significant SNPs: 34 of which were associated with 29 gene candidates (*CG11251*, *goe*, *cpo*, *Mgat1*, and *Ugt86De* were each associated with 2 significant SNPs) either being located within the gene region or nearby and 9 of which were not near any known protein-coding gene (Table 1, Supplementary Table 1, Supplementary Dataset 3). Most gene-associated SNPs were intronic, upstream, or synonymous coding changes, consistent with effects primarily on gene ex^50^pression rather than protein sequence – consistent with the extensive regulatory variation documented in the DGRP. However, *Ugt86De* did contain a SNP that resulted in a nonsynonymous amino acid change that could potentially alter its function. Only one SNP, associated with *Egfr*, passed a Bonferroni-like threshold of 1 divided by the total number of SNPs tested (1/ 1,837,106 yielding a *P*-value of 5.44 x 10^-7^) (Figure 2).

**Figure 2.**
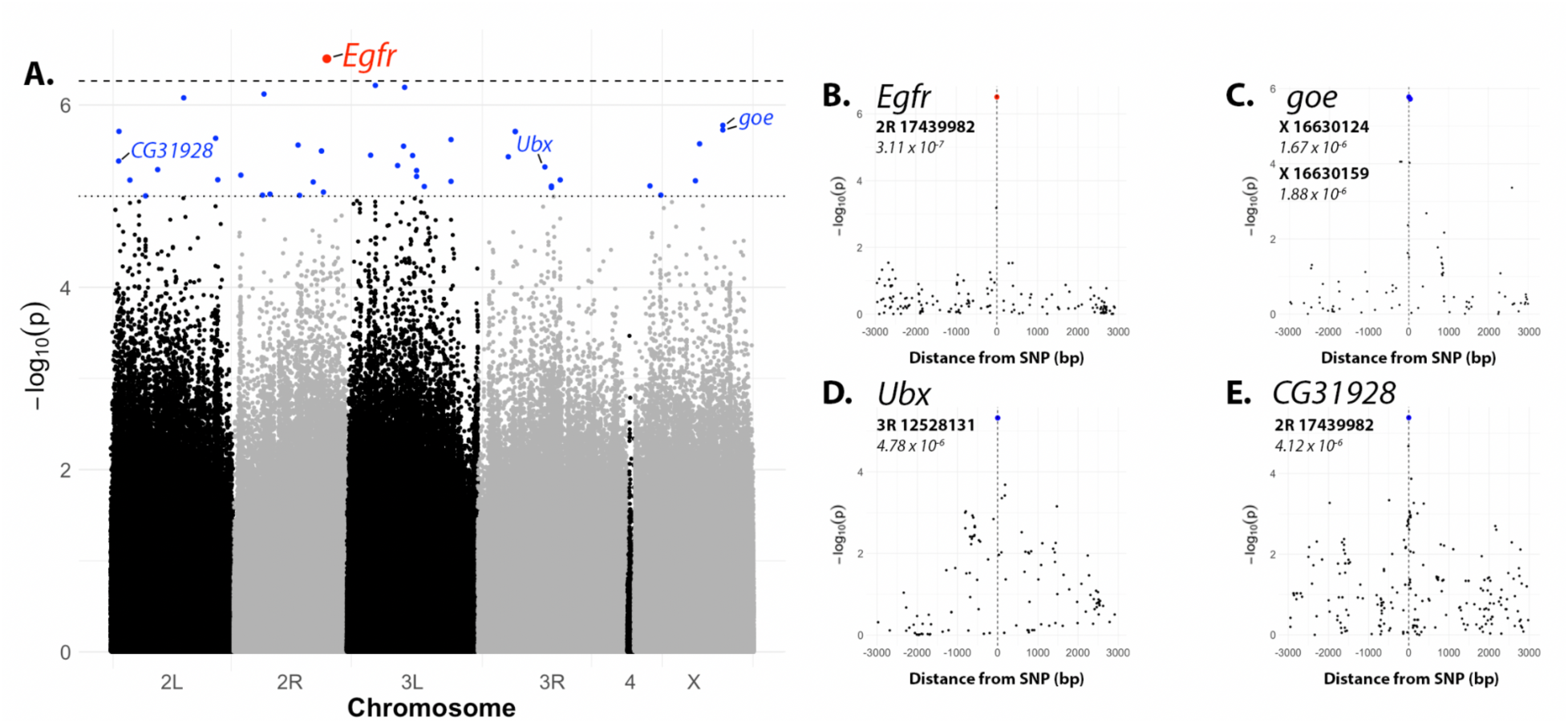
Manhattan plot from GWAS on female timing of mating plug ejection reveals gene candidates. Dotted line – *P*-value threshold of *P* < 1 x 10^-5^, Dashed line – Bonferroni-like corrected *P*-value of 5.44 x 10^-7^ (1/total SNPs in GWAS). Forty-three total significant SNPs pass suggestive *P*-value of 1 x 10^-5^; One gene candidate, *Egfr* passes the more stringent threshold. B-E) Genomic regions containing SNPs associated with genes of interest. The most significant SNP for each gene candidate is surrounded by nonsignificant SNPs evaluated in the GWAS located within 3,000 bp upstream and downstream. The *P*-value of significant SNPs is below their location. B) SNP 2R: 17439982 (linked to *Egfr*) is the SNP most significantly associated with female mating plug ejection timing. C) Two significant SNPs are associated with *goe*, which is known to attenuate Egfr signaling. Both are implicated in a shared regulatory network controlling germ cell differentiation^64^. D) & E) Although only one SNP each is signficantly associated with *Ubx* and *CG31928*, apparent peaks of sub-threshold SNPs surround these candidates as well as others. Data underlying figure are available in Supplementary Dataset 4.

**Table 1.** Female mating plug ejection timing GWAS candiates and associated neuro- and fertility phenotypes and expression. Additional details on gene functions and significant SNPs not associated with known protein coding genes can be found in Supplementary Table 1. List of significant SNPs are availabel in Supplementary Dataset 4.

| Rank Order | P-value | Gene Name | Gene Position | Brain Expression | Abnormal Neuroanatomy | Ovary Expression | Reproductive Process | Female Fertility Effect |
| --- | --- | --- | --- | --- | --- | --- | --- | --- |
| 1 | 3.11E-07 | Egfr | Intron | Y | Y | Y | Oogenesis | Y |
| 6, 7 | 1.67E-06,<br>1.88E-06 | goe | Intron | Y | NA | Y | Oogenesis | Y |
| 11 | 2.32E-06 | Lar | Intron | Y | Y | Y | Oogenesis | Y |
| 15 | 2.83E-06 | cyri | Intron | Y | NA | Y | Oogenesis | Y |
| 16, 17 | 3.20E-06,<br>3.20E-06 | Mgat1 | Intron | N | Y | Y | NA | NA |
| 29 | 6.64E-06 | Dys | Intron | Y | Y | Y | Oogenesis | NA |
| 31 | 6.78E-06 | CG1703 | Intron | Y | NA | Y | NA | NA |
| 34 | 7.72E-06 | kirre | Intron | Y | Y | Y | Oogenesis | Y |
| 35, 37 | 8.05E-06,<br>7.76E-06 | cpo | Intron | Y | Y | Y | Diapause | Y |
| 40 | 9.73E-06 | rg | Intron | Y | Y | Y | NA | NA |
| 43 | 9.93E-06 | eya | Intron | Y | Y | Y | Oogenesis | Y |
| 21 | 4.12E-06 | CG31928 | Upstream | N | NA | Y | Oogenesis | NA |
| 26 | 5.88E-06 | CG30440 | Intron | N | NA | Y | Oogenesis | NA |
| 36 | 7.84E-06 | bbg | Intron | N | NA | Y | Oogenesis | NA |
| 4 | 7.58E-07 | CG1688 | Intron | Y | NA | N | Ovary Development | NA |
| 8 | 1.95E-06 | haf | Intron | Y | Y | N | NA | NA |
| 14 | 2.75E-06 | Ptp52F | Synonymous Coding | Y | Y | N | NA | NA |
| 19 | 3.59E-06 | Lmx1a | Synonymous Coding | Y | NA | N | Ovary Development | Y |
| 20 | 3.69E-06 | Teh1 | Intron | Y | NA | N | NA | NA |
| 28 | 6.61E-06 | sNPF | Intron | Y | NA | N | Egg retention | NA |
| 41 | 9.74E-06 | brp | Intron | Y | NA | N | NA | NA |
| 5 | 8.35E-07 | CG18507 | Synonymous Coding | N | NA | N | NA | NA |
| 9 | 1.95E-06 | Ugt35E2 | Non-synonymous Coding | N | NA | N | NA | NA |
| 10 | 1.95E-06 | Ugt35E2 | Synonymous Coding | N | NA | N | NA | NA |
| 12 | 2.40E-06 | Cpr76Bc | Intron | N | NA | N | NA | NA |
| 18 | 3.56E-06 | CG7465 | Upstream | N | NA | N | NA | NA |
| 23 | 4.78E-06 | Ubx | Intron | N | Y | N | NA | NA |
| 25 | 5.24E-06 | CG11251 | Upstream | N | NA | N | NA | NA |
| 27 | 6.08E-06 | CG11251 | Upstream | N | NA | N | NA | NA |
| 32 | 6.89E-06 | Cpr76Bb | Non-synonymous Coding | N | NA | N | NA | NA |
| 33 | 7.00E-06 | tn | Intron | N | NA | N | NA | NA |

We applied a STRING analysis, based on experimental data, curated databases, and text-mining, to our list of 29 gene candidate genes (Supplementary Figure 6) ^44^. Surprisingly, six connections were present within our candidate list, more than would be expected by chance (PPI enrichment *P*-value 0.0040) (Supplementary Figure 4). GO analysis (Supplementary Figure 2) showed enrichment for cell periphery (*Lar*, *brp*, *Cpr76Bb*, *Cpr76Bc*, *goe*, *Dys*, *CG1688*, *CG31928*, *Ptp52F*, *kirre*, *Teh1*, *Egfr*) and protein tyrosine phosphatase activity (*eya*, *Lar*, *Ptp52f*) among other categories (Supplementary Figures 2-4)^43^. We confirmed the results of the STRING network connections and GO enrichment results with a deeper literature review to verify functional relationships among candidate genes, particularly those modulating RTK signaling, given that *Egfr* is an RTK protein and a key regulator of development signaling^51^. This analysis confirmed that the gene candidates were enriched for functions in development and for localization in the cell periphery, consistent with candidates functioning in developmental signaling pathways (additional description in Supplementary Results 1). Because many candidate genes had broad developmental functions, we next examined whether more specific functional categories were enriched among candidates.

### GWAS gene candidates are enriched for functions in neurodevelopment and in oogenesis

We used Pangea to test for enrichment of Flybase phenotypes (Supplementary Figure 5)^43^. The FBcv terms for abnormal neuroanatomy (FBcv:0000435; Fold Enrichemnt 2.7537; 11/29 candidates) and abnormal behavior (FBcv:0000387; Fold Enrichemnt 2.85; 10/29 candidates) were the top hits. We used FlyAtlas2 expression data to examine tissue expression pattern for our gene candidates and determined that 20/29 candidates are expressed in the brain and/or ventral nerve cord (VNC; referred to as thoracicoabdominal ganglion in FlyAtlas2), and 12 candidates show enriched expression in the brain and/or ganglion (Supplementary Methods 1)^45^. Furthermore, according to Flybase, loss of function alleles of 11 candidate genes have known abnormal neurodevelopmental phenotypes (Table 1, Supplementary Table 1)^46^.

Given the reproductive context of the phenotype, we revisited candidate gene functions in the context of female reproduction. We noted that many candidate genes are expressed in the ovary or have reported mutant phenotypes causing female sterility and/or oogenesis defects in *Drosophila melanogaster*. To formally test this observation, we performed a Fisher’s exact test for enrichment of fertility-related phenotypes and/or known roles in oogenesis among candidate genes relative to all genes tested in the GWAS. This analysis revealed significant enrichment (*p* = 0.011, odds ratio = 3.47), indicating that candidate genes are overrepresented for functions related to female reproduction. Further review of relevant literature revealed additional reproductive roles not fully captured by Flybase annotations, particularly functions in oogenesis and germline development (Table 1 & Supplementary Table 1), and multiple candidates have documented functions in oogenesis within either germline or somatic ovarian tissues^52^.

It should be noted that the GWAS also identified genes with developmental functions that are broadly acting in tissues in addition to or other than the brain, nervous system, and ovary, as well as several genes for which no function has yet been characterized. Furthermore, the GWAS identified some genes with functions not associated with the brain or ovary, notably *Cpr76Bb* and *Cpr76Bc*, two members of the CPR cuticle protein family. In short, the candidate gene list presented here can be used to generate hypotheses for downstream validation.

### Females without a germline show slower and reduced mating plug ejection

Multiple GWAS hits were associated with ovarian function, with the gene candidates spanning a wide range of oogenic processes. Since this association does not appear to be specific to a particular aspect of oocyte or ovary development, it suggests that the relationship might be between mating plug ejection timing and having a full or functional ovary. Since many gene candidates with oogenesis functions also have functions in other tissues across development, standard gene-by-gene functional tests using mutants are unlikely to be informative as any mutant or knockdown females that survive are likely to be unhealthy, causing indirect effects on mating plug ejection. Therefore, to test the role of a functional ovary on oogenesis we tested for effects of total germline removal, rather than by individual oogenesis genes that we detected in the GWAS.

We compared mating plug ejection times of genetically matched females with or without a germline after mating to standard Canton-S males (Figure 3. Supplementary Dataset 5). We observed that females without a germline ejected the mating plug slower and were less likely to eject the plug within the five hours of observation, relative to control females with a germline. Each replicate showed a significant difference by a Welch’s two-sided *t*-test. To combine the independent replicates, we used a linear mixed-effects model with treatment (Experimental vs Control flies) as a fixed effect and replicate as a random effect to account for variation among replicates. Combined analysis showed a significant delay in mating plug ejection timing for females without a germline (*P* < 0.0001). Estimated mean ejection times were 121.9 min for control females and 204.7 min for germline-less females with an estimated effect size of 82.8±9.2 min (t=9.01). Furthermore, in all three replicates we observed a reduced proportion of germline-less females ejecting the mating plug within 5 hours after mating, significant by Fisher’s exact test. A random-effects meta-analysis across these replicates confirmed this significant reduction in the odds of no ejection (odds ratio = 0.18, 95% CI = 0.09–0.36, *z* = -4.93, *P* < 0.0001), with no evidence of heterogeneity among replicates (*P* = 0.7234). Together, these results suggest that the presence of a functional female germline contributes to the regulation of mating plug ejection timing in females (Figure 3).

**Figure 3.**
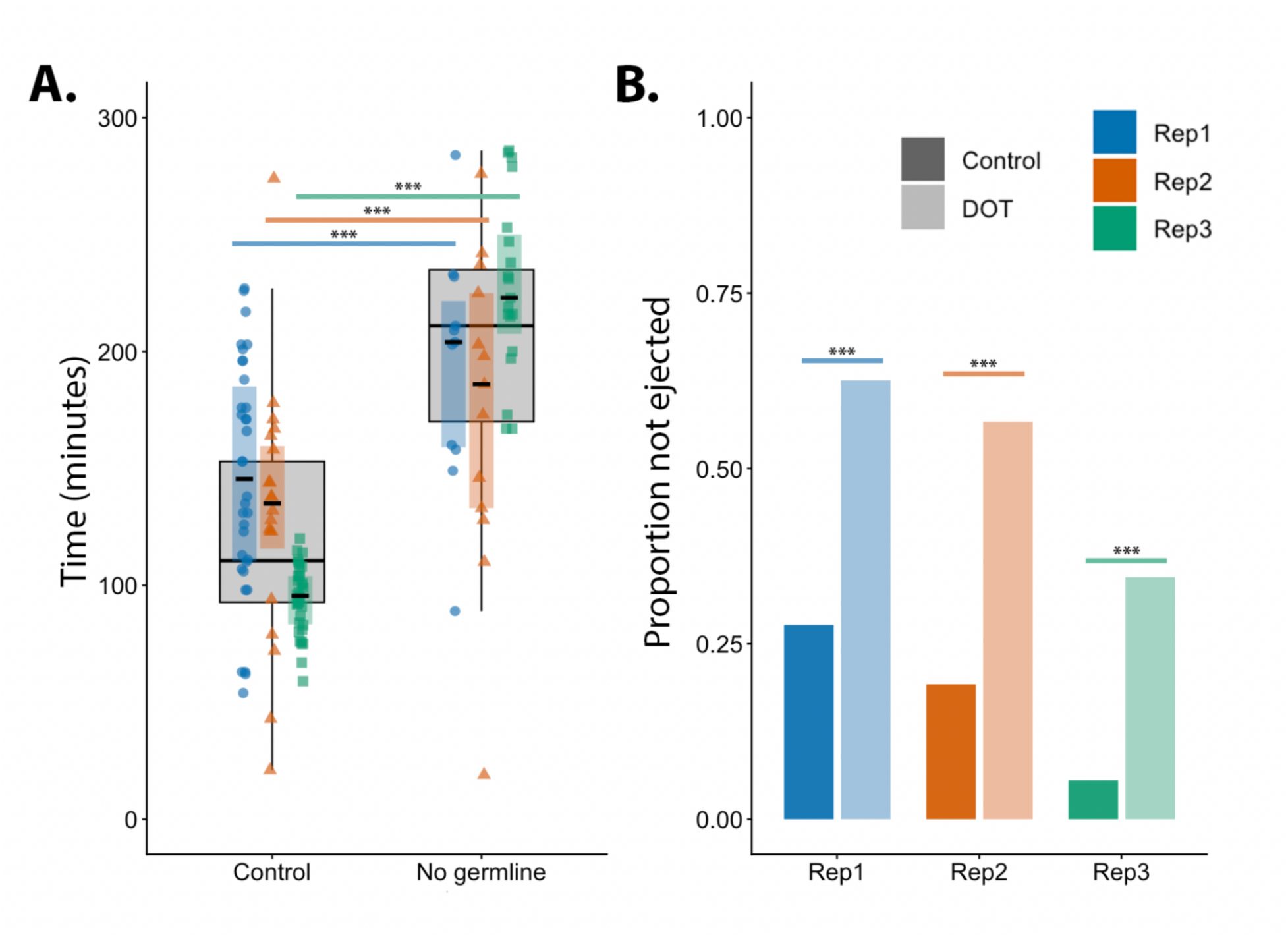
Females without a germline show reduced efficiency of mating plug ejection. Germline-less daughters of females homozygous for the maternal effect mutation *tudor* (DOT; daughter of *tudor*) were compared to genetically matched offspring with a normal germline from mothers heterozygous for *tudor*. A) In three replicates, females without a germline showed increased time to mating plug ejection by Welch’s two-sided *t*-test. Replicates were combined using a linear mixed-effects model, which showed a significant delay in mating plug ejection timing for females without a germline (*P* < 0.0001), with an estimated effect size of 82.8 ± 9.2 min (*t* = 9.01). B) In three replicates, females without a germline were significantly less likely to eject the plug within five hours by a Fisher’s exact test. A random-effects meta-analysis across replicates confirmed this significant reduction in the odds of no ejection (odds ratio = 0.18, 95% CI = 0.09–0.36, *z* = -4.93, *P* < 0.0001). Raw data and data underlying figure are available in Supplementary Dataset 5.

This effect does not appear to be driven by oviposition. The observation chambers used to score mating plug ejection timing in all experiments and DGRP data collection had no food, so females did not lay any eggs, implying that mating plug ejection was not merely a byproduct of oviposition. Additionally, daughters of *tudor* females also ejected the mating plug, despite being unable to lay eggs, further supporting that mating plug ejection is not driven by oviposition. These results suggest that either the presence of a mature ovary accelerates mating plug ejection or that ovary-derived signals promote ejection.

## Discussion

Our study demonstrates that female timing of mating plug ejection, assayed against a fixed male background, exhibits substantial heritable variation within a *D. melanogaster*. This variation likely reflects both female-intrinsic variation and potentially variation in female response to a standard male. Since timing of mating plug ejection is associated with paternity success, this population-level variation can have profound consequences on reproductive outcomes and be subject to selection, including sexual selection. We used the observed phenotypic variation to perform a genome-wide association study on female mating plug ejection timing and identified gene candidates. Functional enrichment analysis revealed that many of the candidate genes are implicated in developmental, neurobiological and reproductive pathways. Using genetic disruption of oogenesis, we demonstrated that rapid, efficient mating plug ejection depends on the presence of a functioning ovary.

### Heritable variation in female mating plug ejection timing

We demonstrated that female timing of mating plug ejection exhibits extensive natural variation within a population of *D. melanogaster*. Differences among the lines were highly repeatable across replicates, and broad-sense heritability was moderate (∼33.7%), comparable to other studies investigating complex behavioral traits with the DGRP^53–55^. These results indicate that genetic differences among females contribute substantially to variation in mating plug ejection timing and suggest that this trait has the potential to evolve under selection. At the same time, the incomplete genetic contribution to the phenotype is consistent with plug ejection being a complex trait shaped by multiple genetic and environmental factors. Since male genotype can contribute to timing of mating plug ejection^33^, we used a standard Canton-S male for all experiments to enable focus on the female-specific genetic variation independent of male genotype. Future studies could investigate correlated male and female effects on mating plug ejection timing across DGRP backgrounds. Correlated or anticorrelated effects could suggest coevolution between the sexes to optimize plug retention and ejection for given combinations of male and female genetic backgrounds, either cooperatively or antagonistically. If these effects were decoupled, this would indicate independently evolving mechanisms regulating this reproductive process.

### Neurodevelopmental and oogenesis genes identified in female mating plug ejection timing GWAS

Genome-wide association studies can identify genes that modulate a phenotype of interest through many diverse mechanisms and pathways simultaneously. We looked for relationships and functional enrichment of our candidate genes in order to form hypotheses regarding the mechanism by which variation in these genes may contribute to population-level differences in female mating plug ejection timing. Our gene candidates are enriched for signaling pathways and developmental processes. Notably, some are known members of the receptor tyrosine kinase pathway or have known genetic interactions with each other. Furthermore, 20 of the 29 candidates are known to be involved in development acting in a variety of tissues and developmental times (Supplementary Figure 2). Taken together, these observations indicate that our GWAS identified many gene candidates that are broadly-acting components of developmental signaling pathways. These findings indicate that the timing of female mating plug ejection is a polygenic and, at least in part, a developmentally embedded trait. Candidates of this type could be needed for overall female vitality and/or sensing and muscular systems which contribute to mating plug ejection, rather than representing specific “ejection genes.” Furthermore, GWAS depends on functional sequence differences among lines to identify associations. Although both *Dh44* and *Dh44-R1* had sequence polymorphism, including nonsynonymous differences across tested fly lines, there were not significant associations with MPE timing in the GWAS analysis. This implies that either the variation had no functional effect, or their effects on the phenotype were small enough as to fall below our threshold for detection as significant.

Beyond those with broad developmental roles, some enriched categories of candidate genes were for those that function in neurodevelopment and oogenesis. That many of the neurodevelopmental genes show enriched expression in the brain or VNC in the adult fly could point to a more specific role in modulating female mating plug ejection behavior beyond simply facilitating general neurodevelopment (Table 1, Supplementary Table 1)^45^. Interestingly, many of the candidates with ovarian function and expression also show enriched expression in the brain or VNC and/or known neurodevelopmental functions. Overlaps between genes implicated in gametogenesis and neural functions have previously been reported; for example, a recent DGRP GWAS study on sperm length similarly identified many neurodevelopmental genes^56,57^. This pattern may reflect the shared use of conserved developmental signaling pathways across tissues rather than tissue-specific pathways. Our top candidate EGFR provides an example of a pleiotropic developmental regulator, functioning in nervous system development and multiple stages of oogenesis as well as development in other tissues^58^. Interestingly, two candidates have neuronal roles that influence egg-laying behavior, *cpo*, which regulates diapause, and *sNPF*, a neuropeptide involved in the regulation of egg retention^59,60^.

### The ovaries influence female mating plug ejection

Given that timing of female mating plug ejection can modulate sperm storage and paternity outcomes and that ejection of the plug is necessary to allow eggs to exit the female, identifying many gene candidates with known functions in oogenesis prompted us to investigate a connection between oogenesis and female plug ejection. We validated this connection by demonstrating that females without a germline have reduced mating plug ejection efficiency. An appealing explanation is that a full functioning ovary takes up substantial space in the abdomen, resulting in steric constraints that affect detection of the mating plug and/or physically facilitating its ejection.

Analogous to our finding of oogenesis genes associating with mating plug ejection timing, Szabad et al. (2019)^61^ reported that daughters of females homozygous for the maternal effect mutant (*tm2gs*), which do not develop a germline, can eject mating plugs. Although Szabad et al. did not quantify the proportion of mutant vs control females that ejected the plug nor the time it took we find that both the proportion and time of ejection are dependent on having functional ovaries. Szabad et al. (2019)^61^ also observed no ejection of the mating plug in females homozygous for *iab-4Db*, which disrupts both ovarian development and a Bithorax Complex miRNA that regulates the expression of the transcription factors Ubx, Abd-A, and Abd-B^62,63^; the first of these genes (*Ubx*) was among our top GWAS hits (Figure 2).

In addition to showing a role for the ovary in mating plug ejection timing, our data suggest that female reproductive state also contributes to modulation of mating plug ejection timing. The fact that daughters of *tudor* females ejected the mating plug despite being unable to lay eggs further supports the idea that mating plug ejection is not driven by oviposition. Under this model, mating plug ejection does not strictly depend on oogenesis, but may occur more rapidly in females that are physiologically prepared for oviposition. Such a mechanism would be consistent with enrichment for candidate genes associated with neuronal functions.

## Conclusion

Our study demonstrates that female timing of mating plug ejection is genetically variable and heritable within *D. melanogaster*. Therefore, selection may act on this trait to modulate optimal timing of female plug ejection. GWAS-identified candidate genes were enriched for developmental functions, in particular neurodevelopment and oogenesis. Our detection of oogenesis gene candidates suggested a relationship between the ovary and timing of mating plug ejection, which we confirmed experimentally. This study establishes female mating plug ejection as an evolvable female trait and provides insight into how female reproductive physiology may shape a post-mating behavior that modulates paternity outcomes.

## Acknowledgments

We thank David Cabello with assistance for initial mating plug ejection timing experiments, Dr. Ben Hopkins for providing an stl file for the 3D printed chambers used in experiments and Jamien Shea for assistance in printing chambers. We thank Norene Buhner and Asha Jain for assistance with fly work. We also thank Drs. Prajal Patel and Emily Rivard for helpful comments on the manuscript. Finally, we thank members of the Wolfner and Clark labs for helpful feedback and support.

## Funding

This work was supported by NIH grant R01-HD059060 awarded to MFW and AGC. JAC was supported by NIH post-doctoral fellowship F32-HD111231 and RMJC and BMV were supported by the Nexus Scholar program sponsored by the Cornell College of Arts and Sciences.

## Notes

### Competing Interest Statement

The authors have declared no competing interest.

## References

1. Firman, R. C., Gasparini, C., Manier, M. K. & Pizzari, T. Postmating Female Control: 20 Years of Cryptic Female Choice. Trends Ecol. Evol. 32, 368–382 (2017).

2. Arbuthnott, D., Fedina, T. Y., Pletcher, S. D. & Promislow, D. E. L. Mate choice in fruit flies is rational and adaptive. Nat. Commun. 8, 13953 (2017).

3. Wedekind, C., Seebeck, T., Bettens, F. & Paepke, A. J. MHC-dependent mate preferences in humans. Proc. R. Soc. B Biol. Sci. 260, 245–249 (1995).

4. Wolff, J. O. & Sherman, P. W. Rodent Societies: An Ecological and Evolutionary Perspective. (University of Chicago Press, 2008).

5. Clutton-Brock, T. Sexual Selection in Males and Females. Science 318, 1882–1885 (2007).

6. Jennions, M. D. Female promiscuity and genetic incompatibility. Trends Ecol. Evol. 12, 251–253 (1997).

7. Searcy, W. A. The Evolutionary Effects of Mate Selection. Annu. Rev. Ecol. Syst. 13, 57–85 (1982).

8. Petersen, R. M. et al. Evidence for genetically-based sperm discrimination in the vaginal tract of a primate species. PLOS Biol. 24, e3003699 (2026).

9. Chow, C. Y., Wolfner, M. F. & Clark, A. G. The Genetic Basis for Male × Female Interactions Underlying Variation in Reproductive Phenotypes of Drosophila. Genetics 186, 1355–1365 (2010).

10. Parker, G. A. SPERM COMPETITION AND ITS EVOLUTIONARY CONSEQUENCES IN THE INSECTS. https://onlinelibrary.wiley.com/doi/10.1111/j.1469-185X.1970.tb01176.x.

11. Eberhard, W. G. Female Control: Sexual Selection by Cryptic Female Choice. (Princeton University Press, 1966).

12. Carlisle, J. A. & Swanson, W. J. Molecular mechanisms and evolution of fertilization proteins. J. Exp. Zoolog. B Mol. Dev. Evol. 336, 652–665 (2021).

13. Ramm, S. A. et al. Sperm competition risk drives plasticity in seminal fluid composition. BMC Biol. 13, 87 (2015).

14. Wigby, S. et al. Seminal Fluid Protein Allocation and Male Reproductive Success. Curr. Biol. 19, 751–757 (2009).

15. Anastasio, O. E., Sinclair, C. S. & Pischedda, A. Cryptic male mate choice for high-quality females reduces male postcopulatory success in future matings. Evolution 77, 1396–1407 (2023).

16. Firman, R. C. & Simmons, L. W. Sperm competition risk generates phenotypic plasticity in ovum fertilizability. Proc. R. Soc. B Biol. Sci. 280, 20132097 (2013).

17. Firman, R. C. & Simmons, L. W. Gametic interactions promote inbreeding avoidance in house mice. Ecol. Lett. 18, 937–943 (2015).

18. Orr, T. J. & Zuk, M. Reproductive delays in mammals: an unexplored avenue for post-copulatory sexual selection. Biol. Rev. 89, 889–912 (2014).

19. Ejaculate–female and sperm–female interactions. in Sperm Biology 247–304 (Academic Press, 2009). doi:10.1016/B978-0-12-372568-4.00007-0.

20. Peckenpaugh, B., Yew, J. Y. & Moyle, L. C. Long-sperm precedence and other cryptic female choices in Drosophila melanogaster. Evolution 79, 467–482 (2025).

21. Doubovetzky, N., Kohlmeier, P., Bal, S. & Billeter, J.-C. Cryptic female choice in response to male pheromones in Drosophila melanogaster. Curr. Biol. 34, 4539–4546.e3 (2024).

22. Ala-Honkola, O. & Manier, M. K. Multiple mechanisms of cryptic female choice act on intraspecific male variation in Drosophila simulans. Behav. Ecol. Sociobiol. 70, 519–532 (2016).

23. Lüpold, S., Manier, M. K., Ala-Honkola, O., Belote, J. M. & Pitnick, S. Male Drosophila melanogaster adjust ejaculate size based on female mating status, fecundity, and age. Behav. Ecol. 22, 184–191 (2011).

24. Hopkins, B. R. et al. Divergent allocation of sperm and the seminal proteome along a competition gradient in Drosophila melanogaster. Proc. Natl. Acad. Sci. 116, 17925–17933 (2019).

25. Chow, C. Y., Wolfner, M. F. & Clark, A. G. Large Neurological Component to Genetic Differences Underlying Biased Sperm Use in Drosophila. Genetics 193, 177–185 (2013).

26. Peckenpaugh, B., Yew, J. Y. & Moyle, L. C. Long-sperm precedence and other cryptic female choices in Drosophila melanogaster. Evolution 79, 467–482 (2025).

27. Wigby, S. et al. The Drosophila seminal proteome and its role in postcopulatory sexual selection. Philos. Trans. R. Soc. B Biol. Sci. 375, 20200072 (2020).

28. Avila, F. W., Wong, A., Sitnik, J. L. & Wolfner, M. F. Don’t pull the plug! the Drosophila mating plug preserves fertility. Fly (Austin) 9, 62–67 (2015).

29. Avila, F. W. et al. Retention of Ejaculate by Drosophila melanogaster Females Requires the Male-Derived Mating Plug Protein PEBme. Genetics 200, 1171–1179 (2015).

30. McDonough-Goldstein, C. E., Pitnick, S. & Dorus, S. Drosophila female reproductive glands contribute to mating plug composition and the timing of sperm ejection. Proc. R. Soc. B Biol. Sci. 289, 20212213 (2022).

31. Brown, N. C. et al. The seminal odorant binding protein Obp56g is required for mating plug formation and male fertility in Drosophila melanogaster. eLife 12, e86409.

32. Bretman, A., Lawniczak, M. K. N., Boone, J. & Chapman, T. A mating plug protein reduces early female remating in *Drosophila melanogaster*. J. Insect Physiol. 56, 107–113 (2010).

33. Lüpold, S. et al. Female mediation of competitive fertilization success in Drosophila melanogaster. Proc. Natl. Acad. Sci. 110, 10693–10698 (2013).

34. Snook, R. R. & Hosken, D. J. Sperm death and dumping in Drosophila. Nature 428, 939–941 (2004).

35. Manier, M. K. et al. Postcopulatory Sexual Selection Generates Speciation Phenotypes in Drosophila. Curr. Biol. 23, 1853–1862 (2013).

36. Laturney, M. & Billeter, J.-C. Drosophila melanogaster females restore their attractiveness after mating by removing male anti-aphrodisiac pheromones. Nat. Commun. 7, 12322 (2016).

37. Lee, K.-M. et al. A Neuronal Pathway that Controls Sperm Ejection and Storage in Female Drosophila. Curr. Biol. 25, 790–797 (2015).

38. Yun, M. et al. Male cuticular pheromones stimulate removal of the mating plug and promote re-mating through pC1 neurons in Drosophila females. eLife 13, RP96013 (2024).

39. Mackay, T. F. C. et al. The Drosophila melanogaster Genetic Reference Panel. Nature 482, 173–178 (2012).

40. Boswell’, E. & Mahowaldt, R. fudor, A Gene Required for Assembly of the Germ Plasm in Drosophila melanogaster.

41. Hopkins, B. R. et al. BMP signaling inhibition in Drosophila secondary cells remodels the seminal proteome and self and rival ejaculate functions. Proc. Natl. Acad. Sci. U. S. A. 116, 24719–24728 (2019).

42. Huang, W. et al. Natural variation in genome architecture among 205 Drosophila melanogaster Genetic Reference Panel lines. Genome Res. 24, 1193–1208 (2014).

43. Hu, Y. et al. PANGEA: a new gene set enrichment tool for Drosophila and common research organisms. Nucleic Acids Res. 51, W419–W426 (2023).

44. Szklarczyk, D. et al. The STRING database in 2023: protein–protein association networks and functional enrichment analyses for any sequenced genome of interest. Nucleic Acids Res. 51, D638–D646 (2023).

45. Krause, S. A., Overend, G., Dow, J. A. T. & Leader, D. P. FlyAtlas 2 in 2022: enhancements to the Drosophila melanogaster expression atlas. Nucleic Acids Res. 50, D1010–D1015 (2022).

46. Öztürk-Çolak, A. et al. FlyBase: updates to the Drosophila genes and genomes database. Genetics 227, iyad211 (2024).

47. Kuznetsova, A., Brockhoff, P. B. & Christensen, R. H. B. lmerTest Package: Tests in Linear Mixed Effects Models. J. Stat. Softw. 82, 1–26 (2017).

48. Bates, D., Mächler, M., Bolker, B. & Walker, S. Fitting Linear Mixed-Effects Models Using lme4. J. Stat. Softw. 67, 1–48 (2015).

49. Viechtbauer, W. Conducting Meta-Analyses in R with the metafor Package. J. Stat. Softw. 36, 1–48 (2010).

50. Huang, W. et al. Genetic basis of transcriptome diversity in Drosophila melanogaster. Proc. Natl. Acad. Sci. 112, E6010–E6019 (2015).

51. Mele, S. & Johnson, T. K. Receptor Tyrosine Kinases in Development: Insights from Drosophila. Int. J. Mol. Sci. 21, 188 (2020).

52. Lmx1a is required for the development of the ovarian stem cell niche in Drosophila | Development | The Company of Biologists. https://journals.biologists.com/dev/article/145/8/dev163394/19364/Lmx1a-is-required-for-the-development-of-the?guestAccessKey=.

53. Arya, G. H. et al. The Genetic Basis for Variation in Olfactory Behavior in Drosophila melanogaster. Chem. Senses 40, 233–243 (2015).

54. Harbison, S. T., McCoy, L. J. & Mackay, T. F. Genome-wide association study of sleep in Drosophila melanogaster. BMC Genomics 14, 281 (2013).

55. Rohde, P. D. et al. Functional Validation of Candidate Genes Detected by Genomic Feature Models. G3 GenesGenomesGenetics 8, 1659–1668 (2018).

56. Ruohola, H. et al. Role of neurogenic genes in establishment of follicle cell fate and oocyte polarity during oogenesis in Drosophila. Cell 66, 433–449 (1991).

57. Syed, Z. A. et al. Genomics of a sexually selected sperm ornament and female preference in Drosophila. *Nat*. Ecol. Evol. 9, 336–348 (2025).

58. Nilson, L. A. & Schüpbach, T. EGF receptor signaling in Drosophila oogenesis. Curr. Top. Dev. Biol. 44, 203–243 (1999).

59. Grmai, L., Michaca, M., Lackner, E., Nampoothiri V.P., N. & Vasudevan, D. Integrated stress response signaling acts as a metabolic sensor in fat tissues to regulate oocyte maturation and ovulation. Cell Rep. 43, 113863 (2024).

60. Schmidt, P. S. et al. An amino acid polymorphism in the couch potato gene forms the basis for climatic adaptation in Drosophila melanogaster. Proc. Natl. Acad. Sci. 105, 16207–16211 (2008).

61. Szabad, J., Peng, J. & Kubli, E. Control of mating plug expelling and sperm storage in Drosophila: A gynandromorph- and mutation-based dissection. Biol. Futura 70, 301–311 (2019).

62. Bender, W. MicroRNAs in the Drosophila bithorax complex. Genes Dev. 22, 14–19 (2008).

63. Ronshaugen, M., Biemar, F., Piel, J., Levine, M. & Lai, E. C. The Drosophila microRNA iab-4 causes a dominant homeotic transformation of halteres to wings. Genes Dev. 19, 2947–2952 (2005).

64. Matsuoka, S., Gupta, S., Suzuki, E., Hiromi, Y. & Asaoka, M. gone early, a Novel Germline Factor, Ensures the Proper Size of the Stem Cell Precursor Pool in the Drosophila Ovary. PLoS ONE 9, e113423 (2014).

