## Supplementary File 1 for "Female genetic variation controlling timing of mating plug ejection in *Drosophila melanogaster*"

### Contents:

**Supplementary Figure 1.** Visual representation of assay for measuring mating plug ejection timing.
**Supplementary Figure 2.** GO Biological Processes Analysis
**Supplementary Figure 3.** GO Cellular Component Analysis
**Supplementary Figure 4.** GO Molecular Function Analysis
**Supplementary Figure 5.** Pangea FlyBase Phenotype Enrichment
**Supplementary Figure 6.** STRING network analysis of gene candidate from GWAS
**Supplementary Figure 7.** Pearson's rank correlation of DGRP line replicates.
**Supplementary Methods 1.** Gene Ontology, Network, and Expression Analysis of Gene Candidates
**Supplementary Results 1.** STRING Analysis and literature summary of gene interactions.

### Additional Associated Files:

**Tables:**
**Supplementary Table 1.** Female mating plug ejection timing GWAS candiates and associated neuro- and fertility phenotypes and expression – Additional Information.

**Datasets:**
**Supplementary Dataset 1.** Raw data for DGRP females.
**Supplementary Dataset 2.** Parametric survival regression model variables and median female mating plug ejection time per DGRP line and replicate
**Supplementary Dataset 3.** Significant SNPs from GWAS
**Supplementary Dataset 4.** Significance scores of SNPs underlying Manhattan Plot
**Supplementary Dataset 5**. Experimental Data from Figure 3

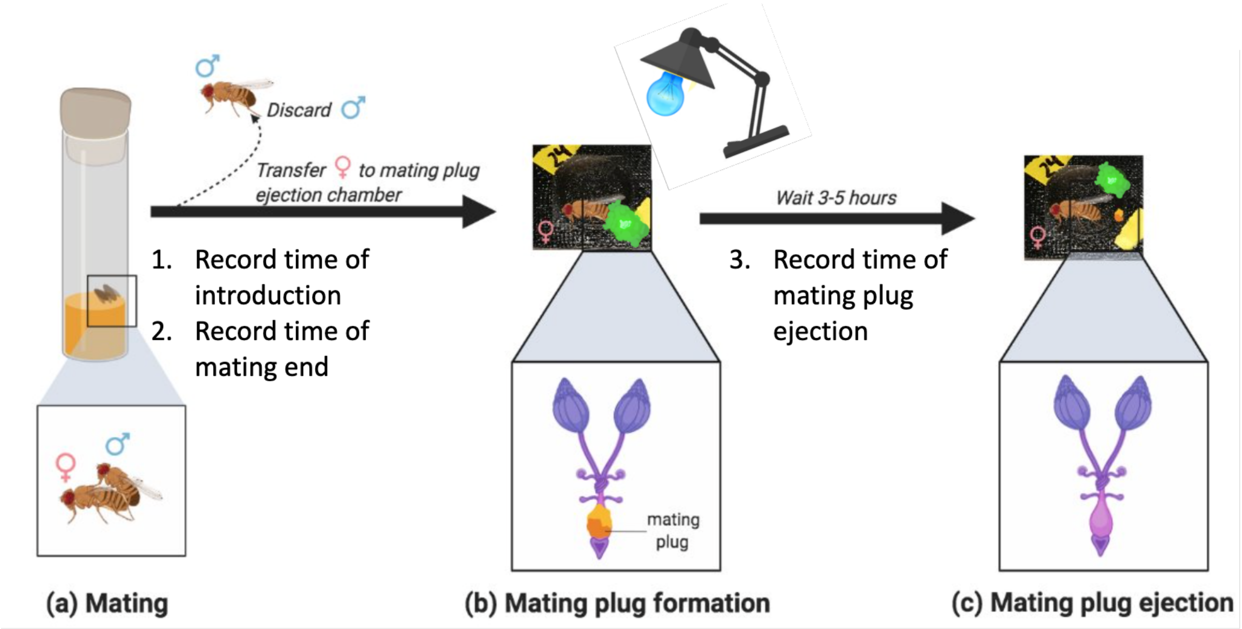


**Supplementary Figure 1. Visual representation of assay for measuring mating plug ejection timing.**

| [Select] | Gene Set Category | Gene Set ID | Gene Set Name | Gene Set Size | Count Overlap Gene | Fold Enrichment | log2 fold | P value | Benjamini & Yekutieli | Overlaping Gene Symbols |
|---|---|---|---|---|---|---|---|---|---|---|
| ▢ | GO Biological Processes | GO:0032502 | developmental process | 3189 | 20 | 3.0207 | 1.595 | 1.587e-7 | 1.449e-3 | sNPF; bbg; cpo; Cpr76Bb; Ptp52F; Egfr; Cpr76Bc; tn; Lar; rg; Lmx1a; Ubx; kirre; haf; eya; Dys; Mgat1; goe; brp; CG32066 |
| ▢ | GO Biological Processes | GO:0048856 | anatomical structure development | 2971 | 18 | 2.9181 | 1.545 | 2.270e-6 | 7.128e-3 | brp; Ptp52F; Cpr76Bb; tn; Cpr76Bc; Lmx1a; Egfr; rg; haf; Lar; kirre; Ubx; Dys; goe; eya; CG32066; Mgat1; bbg |
| ▢ | GO Biological Processes | GO:0042675 | compound eye cone cell differentiation | 15 | 3 | 96.331 | 6.59 | 3.600e-6 | 7.128e-3 | Egfr; rg; eya |
| ▢ | GO Biological Processes | GO:0032501 | multicellular organismal process | 3441 | 19 | 2.6595 | 1.411 | 3.775e-6 | 7.128e-3 | Lar; Cpr76Bc; CG32066; Ubx; kirre; eya; bbg; Mgat1; Ptp52F; brp; Dys; sNPF; tn; cpo; rg; Lmx1a; Egfr; Cpr76Bb; haf |
| ▢ | GO Biological Processes | GO:0030154 | cell differentiation | 1828 | 14 | 3.6888 | 1.883 | 4.652e-6 | 7.128e-3 | goe; tn; Lmx1a; Dys; bbg; kirre; Egfr; Ptp52F; Lar; rg; Ubx; haf; eya; CG32066 |
| ▢ | GO Biological Processes | GO:0048869 | cellular developmental process | 1829 | 14 | 3.6868 | 1.882 | 4.683e-6 | 7.128e-3 | Lmx1a; Dys; bbg; kirre; Egfr; Ptp52F; Lar; rg; Ubx; haf; eya; CG32066; goe; tn |
| ▢ | GO Biological Processes | GO:0007275 | multicellular organism development | 2426 | 15 | 2.9781 | 1.574 | 2.521e-5 | 3.289e-2 | Cpr76Bb; rg; Cpr76Bc; haf; Dys; kirre; Egfr; Ubx; Mgat1; Lar; bbg; eya; brp; Lmx1a; Ptp52F |
| ▢ | GO Biological Processes | GO:0009653 | anatomical structure morphogenesis | 1636 | 12 | 3.5329 | 1.821 | 4.892e-5 | 5.567e-2 | Ubx; rg; CG32066; haf; Egfr; tn; Lar; eya; Dys; bbg; kirre; Ptp52F |
| ▢ | GO Biological Processes | GO:0050793 | regulation of developmental process | 700 | 8 | 5.5046 | 2.461 | 6.414e-5 | 5.857e-2 | cpo; Egfr; goe; Ubx; Lar; kirre; eya; sNPF |
| ▢ | GO Biological Processes | GO:0042692 | muscle cell differentiation | 124 | 4 | 15.5373 | 3.958 | 1.183e-4 | 8.313e-2 | kirre; Ubx; tn; Dys |

**Supplementary Figure 2. GO Biological Processes Analysis**

| [Select] | Gene Set Category | Gene Set ID | Gene Set Name | Gene Set Size | Count Overlap Gene | Fold Enrichment | log2 fold | P value | Benjamini & Yekutieli | Overlaping Gene Symbols |
|---|---|---|---|---|---|---|---|---|---|---|
| ☐ | GO Cellular Component | GO:0071944 | cell periphery | 2056 | 12 | 2.8112 | 1.491 | 4.512e-4 | 1.316e-1 | Lar; brp; Cpr76Bb; Cpr76Bc; goe; Dys; CG1688; CG31928; Ptp52F; kirre; Teh1; Egfr |
| ☐ | GO Cellular Component | GO:0042734 | presynaptic membrane | 35 | 2 | 27.5232 | 4.783 | 2.373e-3 | 2.104e-1 | kirre; brp |
| ☐ | GO Cellular Component | GO:0030863 | cortical cytoskeleton | 38 | 2 | 25.3503 | 4.664 | 2.794e-3 | 2.298e-1 | Dys; brp |
| ☐ | GO Cellular Component | GO:0030054 | cell junction | 481 | 5 | 5.0068 | 2.324 | 2.841e-3 | 2.316e-1 | kirre; Lar; brp; Dys; bbg |
| ☐ | GO Cellular Component | GO:0042995 | cell projection | 748 | 6 | 3.8635 | 1.95 | 3.807e-3 | 2.895e-1 | CG32066; Dys; Egfr; Lar; sNPF; brp |
| ☐ | GO Cellular Component | GO:0030018 | Z disc | 48 | 2 | 20.069 | 4.327 | 4.425e-3 | 2.983e-1 | tn; Dys |
| ☐ | GO Cellular Component | GO:0031674 | I band | 49 | 2 | 19.6594 | 4.297 | 4.608e-3 | 2.993e-1 | Dys; tn |
| ☐ | GO Cellular Component | GO:0070161 | anchoring junction | 165 | 3 | 8.7574 | 3.131 | 4.720e-3 | 2.993e-1 | kirre; bbg; Lar |
| ☐ | GO Cellular Component | GO:0005886 | plasma membrane | 1645 | 9 | 2.6352 | 1.398 | 4.738e-3 | 2.993e-1 | Lar; kirre; brp; Egfr; goe; CG1688; Ptp52F; Teh1; Dys |
| ☐ | GO Cellular Component | GO:0009925 | basal plasma membrane | 58 | 2 | 16.6088 | 4.054 | 6.402e-3 | 3.303e-1 | Lar; Egfr |

**Supplementary Figure 3. GO Cellular Component Analysis**

| [Select] | Gene Set Category | Gene Set ID | Gene Set Name | Gene Set Size | Count Overlap Gene | Fold Enrichment | log2 fold | P value | Benjamini & Yekutieli | Overlaping Gene Symbols |
|---|---|---|---|---|---|---|---|---|---|---|
| ☐ | GO Molecular Function | GO:0004725 | protein tyrosine phosphatase activity | 36 | 3 | 40.1379 | 5.327 | 5.486e-5 | 5.567e-2 | eya; Lar; Ptp52F |
| ☐ | GO Molecular Function | GO:0005001 | transmembrane receptor protein tyrosine phosphatase activity | 7 | 2 | 137.6158 | 7.105 | 8.684e-5 | 6.609e-2 | Lar; Ptp52F |
| ☐ | GO Molecular Function | GO:0019198 | transmembrane receptor protein phosphatase activity | 7 | 2 | 137.6158 | 7.105 | 8.684e-5 | 6.609e-2 | Lar; Ptp52F |
| ☐ | GO Molecular Function | GO:0019904 | protein domain specific binding | 72 | 3 | 20.069 | 4.327 | 4.358e-4 | 1.316e-1 | Lar; Ubx; Dys |
| ☐ | GO Molecular Function | GO:0004721 | phosphoprotein phosphatase activity | 94 | 3 | 15.372 | 3.942 | 9.500e-4 | 1.701e-1 | eya; Lar; Ptp52F |
| ☐ | GO Molecular Function | GO:0016791 | phosphatase activity | 190 | 3 | 7.6051 | 2.927 | 6.979e-3 | 3.502e-1 | eya; Lar; Ptp52F |
| ☐ | GO Molecular Function | GO:0140096 | catalytic activity, acting on a protein | 1513 | 8 | 2.5468 | 1.349 | 9.955e-3 | 4.035e-1 | CG31928; tn; Lar; Ptp52F; Mgat1; eya; Egfr; goe |
| ☐ | GO Molecular Function | GO:0042578 | phosphoric ester hydrolase activity | 224 | 3 | 6.4507 | 2.689 | 1.094e-2 | 4.144e-1 | eya; Lar; Ptp52F |
| ☐ | GO Molecular Function | GO:0038023 | signaling receptor activity | 446 | 4 | 4.3198 | 2.111 | 1.294e-2 | 4.377e-1 | Egfr; Lar; Ptp52F; kirre |
| ☐ | GO Molecular Function | GO:0060089 | molecular transducer activity | 446 | 4 | 4.3198 | 2.111 | 1.294e-2 | 4.377e-1 | Egfr; Lar; Ptp52F; kirre |

**Supplementary Figure 4. GO Molecular Function Analysis**

| [Select] | Gene Set Category | Gene Set ID | Gene Set Name | Gene Set Size | Count Overlap Gene | Fold Enrichment | log2 fold | P value | Benjamini & Yekutieli | Overlaping Gene Symbols |
|---|---|---|---|---|---|---|---|---|---|---|
| ☐ | FlyBase phenotype for classical alleles | FBcv:0000435 | abnormal neuroanatomy | 1924 | 11 | 2.7537 | 1.461 | 1.044e-3 | 1.733e-1 | eya; haf; rg; Lar; tn; Egfr; Ubx; Ptp52F; Mgat1; brp; Dys |
| ☐ | FlyBase phenotype for classical alleles | FBcv:0000387 | abnormal behavior | 1690 | 10 | 2.85 | 1.511 | 1.488e-3 | 1.935e-1 | Dys; cpo; eya; tn; Lar; rg; Egfr; sNPF; Mgat1; brp |
| ☐ | FlyBase phenotype for classical alleles | FBcv:0000396 | abnormal eclosion rhythm | 30 | 2 | 32.1103 | 5.005 | 1.746e-3 | 1.935e-1 | eya; rg |
| ☐ | FlyBase phenotype for classical alleles | FBcv:0000408 | abnormal stress response | 1222 | 8 | 3.1532 | 1.657 | 2.714e-3 | 2.266e-1 | haf; cpo; Lar; Egfr; sNPF; Mgat1; brp; Dys |
| ☐ | FlyBase phenotype for classical alleles | FBcv:0000394 | abnormal circadian rhythm | 286 | 4 | 6.7364 | 2.752 | 2.730e-3 | 2.266e-1 | Lar; brp; rg; eya |
| ☐ | FlyBase phenotype for classical alleles | FBcv:0000428 | abnormal cell size | 331 | 4 | 5.8206 | 2.541 | 4.606e-3 | 2.993e-1 | sNPF; tn; Egfr; Ubx |
| ☐ | FlyBase phenotype for classical alleles | FBcv:0000392 | hyperactive | 59 | 2 | 16.3273 | 4.029 | 6.618e-3 | 3.376e-1 | Egfr; rg |
| ☐ | FlyBase phenotype for classical alleles | FBcv:0000670 | abnormal eclosion | 65 | 2 | 14.8202 | 3.889 | 7.984e-3 | 3.567e-1 | eya; rg |
| ☐ | FlyBase phenotype for classical alleles | FBcv:0000414 | abnormal locomotor behavior | 1157 | 7 | 2.9141 | 1.543 | 8.106e-3 | 3.567e-1 | brp; Dys; cpo; tn; rg; sNPF; Mgat1 |
| ☐ | FlyBase phenotype for classical alleles | FBcv:0000679 | abnormal circadian behavior | 212 | 3 | 6.8159 | 2.769 | 9.419e-3 | 3.910e-1 | eya; brp; rg |

**Supplementary Figure 5. Pangea FlyBase Phenotype Enrichment**

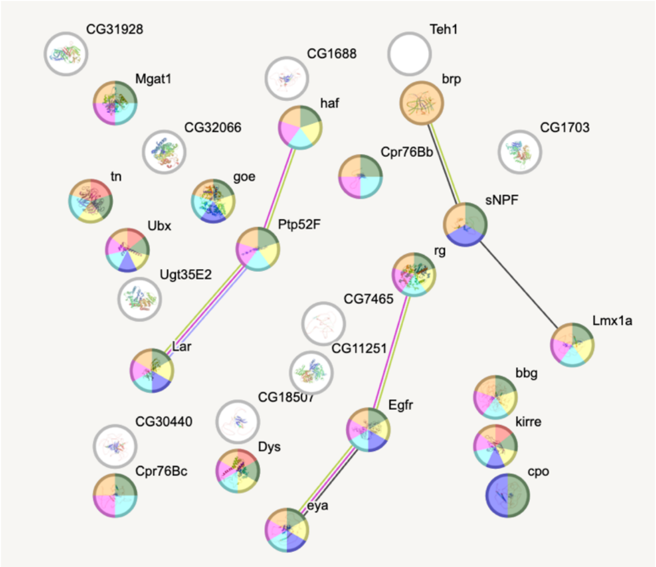


**Network Stats**

| | |
|---|---|
| number of nodes: | 29 |
| number of edges: | 6 |
| average node degree: | 0.414 |
| avg. local clustering coefficient: | 0.207 |
| expected number of edges: | 1 |
| PPI enrichment p-value: | 0.00396 |

*your network has significantly more interactions than expected (what does that mean?)*

**Functional enrichments in your network**

*explain columns*

**Biological Process (Gene Ontology)**

| GO-term | description | count in network | strength | signal | false discovery rate |
|---|---|---|---|---|---|
| GO:0042675 | Compound eye cone cell differentiation | 3 of 12 | 2.08 | 0.77 | 0.0127 |
| GO:0042692 | Muscle cell differentiation | 4 of 101 | 1.28 | 0.48 | 0.0486 |
| GO:0032502 | Developmental process | 18 of 2666 | 0.51 | 0.43 | 0.0033 |
| GO:0030154 | Cell differentiation | 13 of 1558 | 0.6 | 0.42 | 0.0127 |
| GO:0050793 | Regulation of developmental process | 8 of 673 | 0.76 | 0.39 | 0.0465 |
| GO:0048856 | Anatomical structure development | 16 of 2531 | 0.48 | 0.37 | 0.0127 |
| GO:0007275 | Multicellular organism development | 14 of 2125 | 0.5 | 0.34 | 0.0298 |
| GO:0032501 | Multicellular organismal process | 18 of 3342 | 0.41 | 0.32 | 0.0170 |

*(less ...)*

**Supplementary Figure 6. STRING network analysis of gene candidate from GWAS**

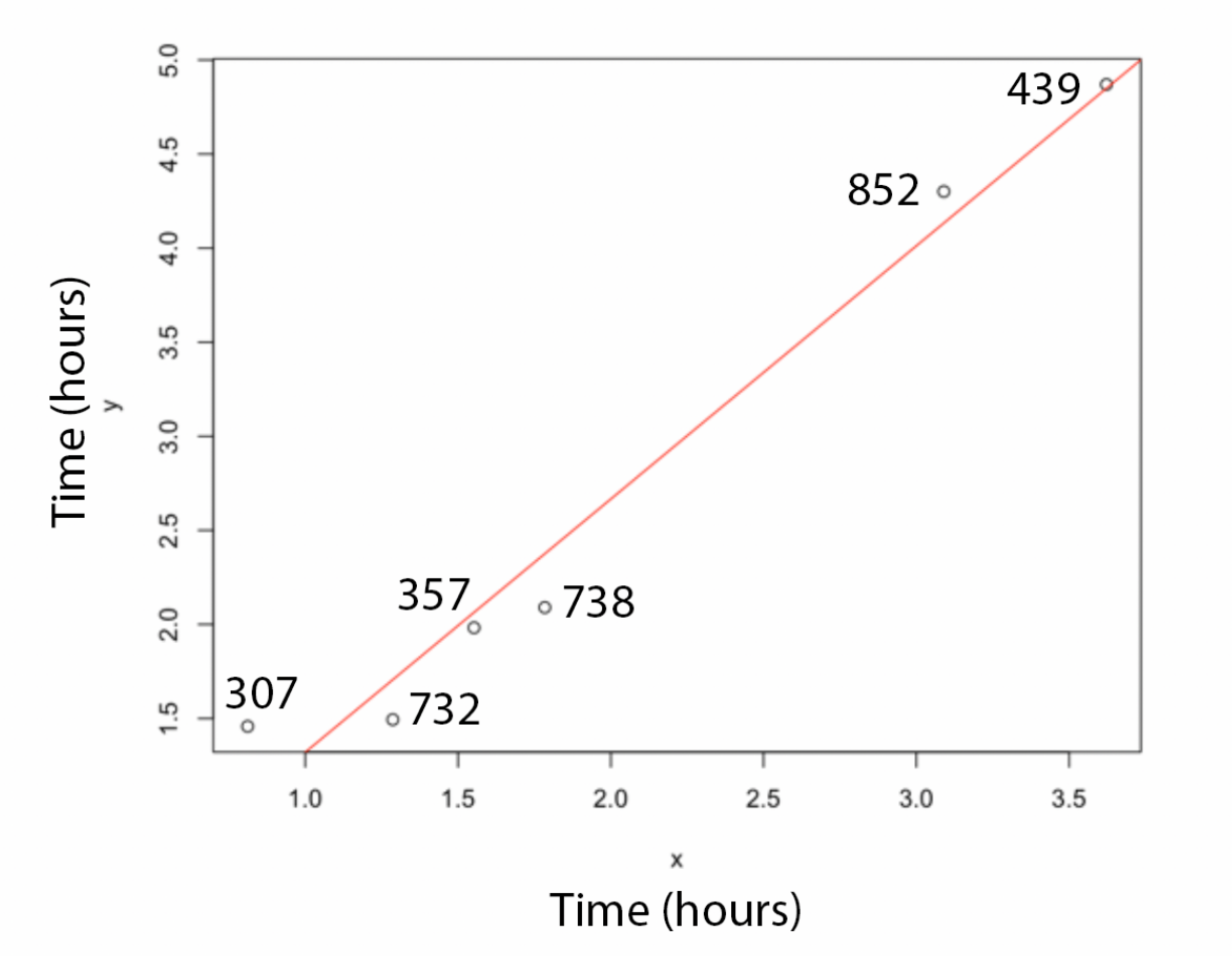


**Supplementary Figure 7.** Pearson's rank correlation of DGRP line replicates.

**Supplementary Methods 1.**

**Gene Ontology, Network, and Expression Analysis of Gene Candidates**

*Enrichment of Gene Function.* Pangea was used to perform gene ontology enrichment analysis. Pangea was used to investigate gene ontology and phenotypic enrichment analysis. Using GO Hierarchy, we investigated GO enrichment for Biological Processes, Cellular Components, and Molecular Functions (Supplementary Figure 2-4) of our 29 Candidates.

We also investigated whether our gene candidate list was enriched for Fly Base phenotypes associated with classic alleles (Fbcv) using Pangea (Supplementary Figure 5). We also tested for enrichment for functions associated with female fertility and oogenesis within our candidate list. From Flybase we downloaded a list of all D. melanogaster genes with known functions in oogenesis or that have classic alleles causing female sterility or semi-sterility phenotypes. We collated these lists and removed duplicates (2173 genes). We then determined how many of the genes examined in our GWAS or our gene candidate list were found on this list of genes or not. Using these values, we performed a Fisher's exact test to determine if our candidate list was enriched for genes effecting the oogenesis or female fertility more broadly. Finally, we did a literature search for our gene list to identify known functions in reproduction which we summarized in Table 1 with additional details on supporting references in Supplementary Table 1. The results of this enrichment analysis are likely conservative since not all genes examined in our GWAS have deep SNP coverage.

*STRING Network.* We input our list of 29 gene candidates into STRING using the D. melanogaster database. We generated a full STRING network using the active interaction sources of Textmining, Experiments, and Databases (Supplementary Figure 6).

*Tissue-Specific Expression*. We investigated tissue-specific expression using data from FlyAtlas 2. Gene-level FPKM values were log2-transformed and we calculated tissue enrichment for each gene and the difference between expression in the focal tissue (Ovary, Brain, or Thoracicoabdominal Ganglion) and the mean expression across all other female tissues. Genes with >2-fold higher expression relative to other tissues were considered to have enriched expression in the focal tissue.
