## Supplementary Table 1 for "Female genetic variation controlling timing of mating plug ejection in *Drosophila melanogaster*": Supplementary_Full_Table_wReferences.docx

| **Rank Order** | **P-value** | **Flybase ID** | **Gene Name** | **Chr Position** | **Gene Position** | **Distance from Gene** | **Expressed in Brain**^1^ | **Enriched Expression in Brain** | **Abnormal Neuroanatomy**^2^ | **Ovary Expression**^1^ | **Enriched Ovary Expression** | **Reproductive Process** | **Somatic or Germline** | **Function** | **Egg Devo Stage** | **Female Fertility Effect** | **Male Fertility Effect?** |
| --- | --- | --- | --- | --- | --- | --- | --- | --- | --- | --- | --- | --- | --- | --- | --- | --- | --- |
| 1 | 3.109E-07 | FBgn0003731 | Egfr | 2R_17439982_SNP | Intron | 0 | Y | N | Y | Y | N | Oogenesis^3–7^ | Somatic | Egg patterning and Egg chamber organization | Early to Late | Y | Y^8^ |
| 6, 7 | 1.67E-06, 1.88E-06 | FBgn0027528 | goe | X_16630124_SNP | Intron | 0 | Y | N | NA | Y | N | Oogenesis^9^ | Germline | PGC differentiation | Early | Y | NA |
| 11 | 2.319E-06 | FBgn0000464 | Lar | 2L_19642047_SNP | Intron | 0 | Y | Y | Y | Y | N | Oogenesis^10–13^ | Both | follicle cell-driven egg elongation | Middle to Late | Y | Y^14^ |
| 15 | 2.829E-06 | FBgn0052066 | cyri | 3L_10639012_SNP | Intron | 0 | Y | N | NA | Y | N | Oogenesis^15^ | Somatic | Border Cell Migration and proper egg formation | Middle to Late | Y | NA |
| 16, 17 | 3.20E-06, 3.20E-06 | FBgn0034521 | Mgat1 | 2R_16447824_SNP | Intron | 0 | N | N | Y | Y | N | NA | NA | NA | NA | NA | Y^16^ |
| 29 | 6.636E-06 | FBgn0260003 | Dys | 3R_15392243_SNP | Intron | 0 | Y | N | Y | Y | N | Oogenesis^17^ | Both | Oocyte Polarity Establishment | Early | NA | NA |
| 31 | 6.781E-06 | FBgn0030321 | CG1703 | X_11507340_SNP | Intron | 0 | Y | N | NA | Y | Y | NA | NA | NA | NA | NA | NA |
| 34 | 7.723E-06 | FBgn0028369 | kirre | X_2998392_SNP | Intron | 0 | Y | Y | Y | Y^18^ | N | Oogenesis^18–21^ | Both | PGC differentiation, ovariole muscle contractions | Early and Late | Y | NA |
| 35, 37 | 8.05E-06, 7.76E-06 | FBgn0263995 | cpo | 3R_13754519_SNP | Intron | 0 | Y | Y | Y | Y | N | Diapause | NA | Diapause arrest | Early | Y | NA |
| 40 | 9.735E-06 | FBgn0266098 | rg | X_5033708_SNP | Intron | 0 | Y | Y | Y | Y | N | NA | NA | NA | NA | NA | NA |
| 43 | 9.930E-06 | FBgn0000320 | eya | 2L_6534815_SNP | Intron | 0 | Y | Y | Y | Y | N | Oogenesis^22^ | Somatic | Egg Chamber Organization | Early to Late | Y | Y^23^ |
| 21 | 4.122E-06 | FBgn0051928 | CG31928 | 2L_1491813_SNP | Upstream | 482 | N | N | NA | Y | Y | Oogenesis^24,25^ | Somatic | Eggshell Structure | Late | NA | NA |
| 26 | 5.882E-06 | FBgn0050440 | CG30440 | 2R_1365028_SNP | Intron | 0 | N | N | NA | Y | Y | Oogenesis^26^ | Somatic | Border Cell migration | Middle | NA | NA |
| 36 | 7.841E-06 | FBgn0087007 | bbg | 3L_14516677_SNP | Intron | 0 | N | N | NA | Y | N | Oogenesis^27^ | Somatic | Border Cell migration | Late | NA | NA |
| 4 | 7.579E-07 | FBgn0027589 | CG1688 | 2R_5693208_SNP | Intron | 0 | Y | Y | NA | N | N | NA | NA | NA | NA | NA | NA |
| 8 | 1.946E-06 | FBgn0261509 | haf | 2L_1557910_INS | Intron | 0 | Y | Y | Y | N | N | NA | NA | NA | NA | NA | NA |
| 14 | 2.751E-06 | FBgn0034085 | Ptp52F | 2R_12060186_SNP | Synonymous Coding | 0 | Y | Y | Y | N | N | NA | NA | NA | NA | NA | NA |
| 19 | 3.585E-06 | FBgn0052105 | Lmx1a | 3L_12328504_SNP | Synonymous Coding | 0 | Y | Y | NA | N | N | Ovary Development^28^ | Somatic | Ovary development | NA | Y | N |
| 20 | 3.695E-06 | FBgn0037766 | Teh1 | 3R_5683836_SNP | Intron | 0 | Y | Y | NA | N | N | NA | NA | NA | NA | NA | NA |
| 28 | 6.611E-06 | FBgn0032840 | sNPF | 2L_20029500_SNP | Intron | 0 | Y | Y | NA | N | N | Egg Retention^29^ | NA | NA | NA | NA | NA |
| 41 | 9.741E-06 | FBgn0259246 | brp | 2R_5418787_SNP | Intron | 0 | Y | Y | NA | N | N | NA | NA | NA | NA | NA | NA |
| 2 | 6.095E-07 | NA | NA | 3L_5369836_SNP | NA | NA | NA | NA | NA | N | NA | NA | NA | NA | NA | NA | NA |
| 3 | 6.403E-07 | NA | NA | 3L_10846322_SNP | NA | NA | NA | NA | NA | N | NA | NA | NA | NA | NA | NA | NA |
| 13 | 2.667E-06 | NA | NA | X_12283844_SNP | NA | NA | NA | NA | NA | N | NA | NA | NA | NA | NA | NA | NA |
| 22 | 4.620E-06 | NA | NA | 3L_9526581_SNP | NA | NA | NA | NA | NA | N | NA | NA | NA | NA | NA | NA | NA |
| 24 | 5.117E-06 | NA | NA | 2L_8799483_SNP | NA | NA | NA | NA | NA | N | NA | NA | NA | NA | NA | NA | NA |
| 30 | 6.664E-06 | NA | NA | 2L_3601165_SNP | NA | NA | NA | NA | NA | N | NA | NA | NA | NA | NA | NA | NA |
| 38 | 9.0346E-06 | NA | NA | 2R_16801854_SNP | NA | NA | NA | NA | NA | N | NA | NA | NA | NA | NA | NA | NA |
| 39 | 9.492E-06 | NA | NA | 2R_6810365_SNP | NA | NA | NA | NA | NA | N | NA | NA | NA | NA | NA | NA | NA |
| 42 | 9.789E-06 | NA | NA | 2R_12342554_SNP | NA | NA | NA | NA | NA | N | NA | NA | NA | NA | NA | NA | NA |
| 5 | 8.347E-07 | FBgn0028527 | CG18507 | 2L_13669209_SNP | Synonymous Coding | 0 | N | N | NA | N | N | NA | NA | NA | NA | NA | NA |
| 9 | 1.953E-06 | FBgn0040255 | Ugt35E2 | 3R_6992655_SNP | Non-synonymous Coding | 0 | N | N | NA | N | N | NA | NA | NA | NA | NA | NA |
| 10 | 1.953E-06 | FBgn0040255 | Ugt35E2 | 3R_6992666_SNP | Synonymous Coding | 0 | N | N | NA | N | N | NA | NA | NA | NA | NA | NA |
| 12 | 2.398E-06 | FBgn0036880 | Cpr76Bc | 3L_19515771_SNP | Intron | 0 | N | N | NA | N | N | NA | NA | NA | NA | NA | NA |
| 18 | 3.558E-06 | FBgn0035551 | CG7465 | 3L_4480003_SNP | Upstream | 280 | N | N | NA | N | N | NA | NA | NA | NA | NA | NA |
| 23 | 4.785E-06 | FBgn0003944 | Ubx | 3R_12528131_SNP | Intron | 0 | N | N | Y | N | N | NA | NA | NA | NA | NA | NA |
| 25 | 5.238E-06 | FBgn0036346 | CG11251 | 3L_13083226_DEL | Upstream | 158 | N | N | NA | N | N | NA | NA | NA | NA | NA | NA |
| 27 | 6.082E-06 | FBgn0036346 | CG11251 | 3L_13083254_SNP | Upstream | 185 | N | N | NA | N | N | NA | NA | NA | NA | NA | NA |
| 32 | 6.892E-06 | FBgn0036879 | Cpr76Bb | 3L_19512864_SNP | Non-synonymous Coding | 0 | N | N | NA | N | N | NA | NA | NA | NA | NA | NA |
| 33 | 6.997E-06 | FBgn0265356 | tn | 2R_14886443_SNP | Intron | 0 | N | N | NA | N | N | NA | NA | NA | NA | NA | NA |

1.	Leader, D. P., Krause, S. A., Pandit, A., Davies, S. A. & Dow, J. A. T. FlyAtlas 2: a new version of the Drosophila melanogaster expression atlas with RNA-Seq, miRNA-Seq and sex-specific data. *Nucleic Acids Res.* **46**, D809–D815 (2018).

2.	Öztürk-Çolak, A. *et al.* FlyBase: updates to the Drosophila genes and genomes database. *Genetics* **227**, iyad211 (2024).

3.	Duchek, P. & Rørth, P. Guidance of cell migration by EGF receptor signaling during Drosophila oogenesis. *Science* **291**, 131–133 (2001).

4.	Gilboa, L. & Lehmann, R. Soma–germline interactions coordinate homeostasis and growth in the Drosophila gonad. *Nature* **443**, 97–100 (2006).

5.	González-Reyes, A., Elliott, H. & St Johnston, D. Polarization of both major body axes in Drosophila by gurken-torpedo signalling. *Nature* **375**, 654–658 (1995).

6.	James, K. E., Dorman, J. B. & Berg, C. A. Mosaic analyses reveal the function of Drosophila Ras in embryonic dorsoventral patterning and dorsal follicle cell morphogenesis. *Development* **129**, 2209–2222 (2002).

7.	Fregoso Lomas, M., Hails, F., Boisclair Lachance, J.-F. & Nilson, L. A. Response to the Dorsal Anterior Gradient of EGFR Signaling in *Drosophila* Oogenesis Is Prepatterned by Earlier Posterior EGFR Activation. *Cell Rep.* **4**, 791–802 (2013).

8.	Kiger, A. A., White-Cooper, H. & Fuller, M. T. Somatic support cells restrict germline stem cell self-renewal and promote differentiation. *Nature* **407**, 750–754 (2000).

9.	Matsuoka, S., Gupta, S., Suzuki, E., Hiromi, Y. & Asaoka, M. gone early, a Novel Germline Factor, Ensures the Proper Size of the Stem Cell Precursor Pool in the Drosophila Ovary. *PLoS ONE* **9**, e113423 (2014).

10.	Fat2 acts through the WAVE regulatory complex to drive collective cell migration during tissue rotation | Journal of Cell Biology | Rockefeller University Press. https://rupress.org/jcb/article/212/5/591/38358/Fat2-acts-through-the-WAVE-regulatory-complex-to.

11.	Krueger, N. X. *et al.* Functions of the Ectodomain and Cytoplasmic Tyrosine Phosphatase Domains of Receptor Protein Tyrosine Phosphatase Dlar In Vivo. *Mol. Cell. Biol.* **23**, 6909–6921 (2003).

12.	Bateman, J., Reddy, R. S., Saito, H. & Van Vactor, D. The receptor tyrosine phosphatase Dlar and integrins organize actin filaments in the *Drosophila* follicular epithelium. *Curr. Biol.* **11**, 1317–1327 (2001).

13.	Frydman, H. M. & Spradling, A. C. The receptor-like tyrosine phosphatase Lar is required for epithelial planar polarity and for axis determination with Drosophila ovarian follicles. *Development* **128**, 3209–3220 (2001).

14.	Srinivasan, S., Mahowald, A. P. & Fuller, M. T. The receptor tyrosine phosphatase Lar regulates adhesion between Drosophila male germline stem cells and the niche. *Development* **139**, 1381–1390 (2012).

15.	CYRI controls epidermal wound closure and cohesion of invasive border cell cluster in Drosophila | Journal of Cell Biology | Rockefeller University Press. https://rupress.org/jcb/article/223/12/e202310153/277044/CYRI-controls-epidermal-wound-closure-and-cohesion.

16.	Sarkar, M. *et al.* Null Mutations in Drosophila N-Acetylglucosaminyltransferase I Produce Defects in Locomotion and a Reduced Life Span *. *J. Biol. Chem.* **281**, 12776–12785 (2006).

17.	Shcherbata, H. R. *et al.* Dissecting muscle and neuronal disorders in a Drosophila model of muscular dystrophy. *EMBO J.* **26**, 481–493 (2007).

18.	Valer, F. B., Machado, M. C. R., Silva-Junior, R. M. P. & Ramos, R. G. P. Expression of Hbs, Kirre, and Rst during Drosophila ovarian development. *genesis* **56**, e23242 (2018).

19.	Ben-Zvi, D. S. & Volk, T. Escort cell encapsulation of Drosophila germline cells is maintained by irre cell recognition module proteins. *Biol. Open* **8**, bio039842 (2019).

20.	Valer, F. B., Machado, M. C. R., Silva-Junior, R. M. P. & Ramos, R. G. P. Expression of Hbs, Kirre, and Rst during Drosophila ovarian development. *genesis* **56**, e23242 (2018).

21.	Lobell, A. S., Kaspari, R. R., Serrano Negron, Y. L. & Harbison, S. T. The Genetic Architecture of Ovariole Number in Drosophila melanogaster: Genes with Major, Quantitative, and Pleiotropic Effects. *G3 GenesGenomesGenetics* **7**, 2391–2403 (2017).

22.	Bai, J. & Montell, D. Eyes absent, a key repressor of polar cell fate during Drosophila oogenesis. *Development* **129**, 5377–5388 (2002).

23.	Fabrizio, J. J., Boyle, M. & DiNardo, S. A somatic role for eyes absent (eya) and sine oculis (so) in Drosophila spermatocyte development. *Dev. Biol.* **258**, 117–128 (2003).

24.	Fakhouri, M. *et al.* Minor proteins and enzymes of the *Drosophila* eggshell matrix. *Dev. Biol.* **293**, 127–141 (2006).

25.	Tootle, T. L., Williams, D., Hubb, A., Frederick, R. & Spradling, A. Drosophila Eggshell Production: Identification of New Genes and Coordination by Pxt. *PLOS ONE* **6**, e19943 (2011).

26.	Kanca, O. Regulation and targets of Mal-D during border cell migration in Drosophila melanogaster oogenesis. https://archiv.ub.uni-heidelberg.de/volltextserver/7953/ (2007) doi:10.11588/heidok.00007953.

27.	Aranjuez, G., Kudlaty, E., Longworth, M. S. & McDonald, J. A. On the Role of PDZ Domain-Encoding Genes in Drosophila Border Cell Migration. *G3 GenesGenomesGenetics* **2**, 1379–1391 (2012).

28.	Allbee, A. W., Rincon-Limas, D. E. & Biteau, B. Lmx1a is required for the development of the ovarian stem cell niche in Drosophila. *Development* **145**, dev163394 (2018).

29.	Grmai, L., Michaca, M., Lackner, E., Nampoothiri V.P., N. & Vasudevan, D. Integrated stress response signaling acts as a metabolic sensor in fat tissues to regulate oocyte maturation and ovulation. *Cell Rep.* **43**, 113863 (2024).
